# Genome-wide A→G and C→T Mutations Induced by Functional TadA Variants in *Escherichia coli*

**DOI:** 10.1101/2024.08.29.610230

**Authors:** Hao Wang, Zhengxin Dong, Jingyi Shi, Lei Chen, Tao Sun, Weiwen Zhang

**Author notes:** To whom all correspondence should be addressed: Prof. Dr. Tao Sun, Center for Biosafety Research and Strategy, Tianjin University, Tianjin 30072, P.R. China., Prof. Dr. Weiwen Zhang, Laboratory of Synthetic Microbiology, School of Chemical Engineering & Technology, Tianjin University, Tianjin 300072, P.R. China.

## Abstract

The fusion expression of DNA replication-related proteins with nucleotide deaminase enzymes promotes random mutations in bacterial genomes, thereby increasing genetic diversity among population. Most previous studies have focused on cytosine deaminase, which produces only C→T mutations, significantly limiting the variety of mutation types. In this study, we developed a fusion expression system by combining DnaG (RNA primase) with adenine deaminase TadA-8e (DnaG-TadA) in *Escherichia coli*, which is capable of rapidly introducing A→G mutations into the *E. coli* genome, resulting in a 664-fold increase in terms of mutation rate. Additionally, we engineered a dual-functional TadA variant, TadAD, and then fused it with DnaG. This construct introduced both C→T and A→G mutations into the *E. coli* genome, with the mutation rate further increased by 370-fold upon co-expression with an uracil glycosylase inhibitor (DnaG-TadAD-UGI). We applied DnaG-TadA and DnaG-TadAD-UGI systems to the adaptive laboratory evolution for Cd^2+^ and kanamycin resistance, achieving an 8.0 mM Cd^2+^ and 200 μg/mL kanamycin tolerance within just 17 days and 132 hours, respectively. Compared to conventional evolution methods, the final tolerance levels were increased by 320% and 266%, respectively. Our work offers a novel strategy for random mutagenesis in *E. coli* and potentially other prokaryotic species.

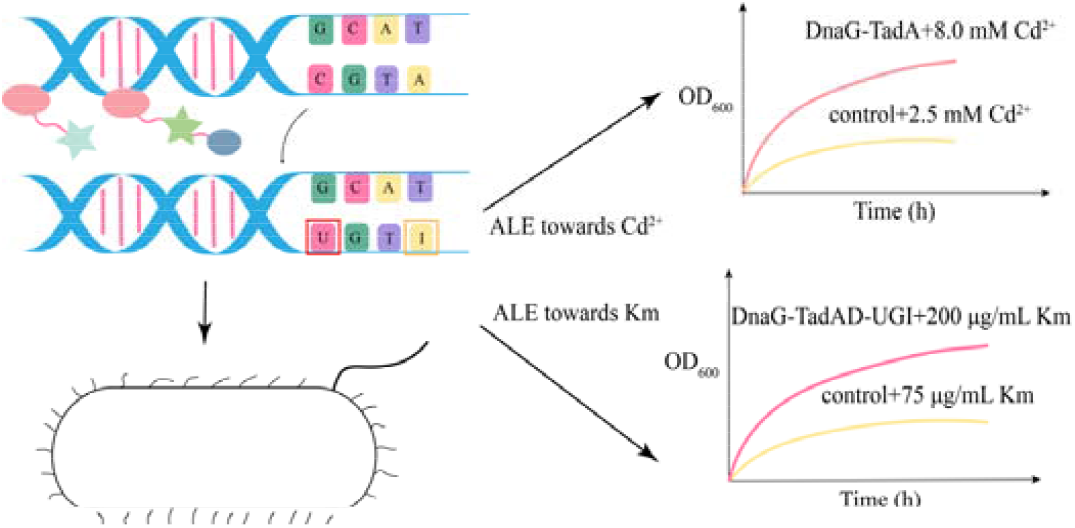

For TOC only

## INTRODUCTION

Random mutations play a crucial role in chassis evolution by introducing genetic diversity into the genome, promoting the emergence of individuals better adapted to environmental changes or capable of producing higher yields of specific compounds^1^. However, the natural rate of random mutations in bacteria severely limits the speed of the laboratory adaptive evolution. To overcome this, physical mutagenesis, chemical mutagenesis, and physical methods such as Atmospheric Pressure Plasma at Room Temperature (ARTP) have been commonly used to increase the random mutation frequency in the past^2-4^. However, they generally pose significant problems with inhibiting cell growth, and the separation of mutation and screening processes can greatly prolong the time needed to identify desirable strains. In recent years, tools such as MP6 (Mutagenesis Plasmids 6) and GREACE (Genome Replication Engineering Assisted Continuous Evolution), developed by disrupting DNA replication fidelity, have significantly increased random mutation rates; however, they also have a considerable impact on cell growth^5, 6^. Base editing^7, 8^ and TRIDENT^9^ (Targeted *In vivo* Diversification Enabled by T7 RNAP) offer innovative methods for introducing mutations into the genome, making significant contributions to genome modification and protein-directed evolution. However, they typically target mutations within a very narrow genomic range. CoMuTER (Confined Mutagenesis using a Type I-E CRISPR-Cas system) combines Cas3 with base editing to introduce mutations within a 55 kb gene region in yeast *Saccharomyces cerevisiae*^10^. However, because Cas3 possesses both helicase and nuclease activities, it has the potential disadvantage to cause large-scale genome deletions^11^.

In recent years, research has been conducted to achieve random base editing of DNA by expressing cytosine deaminase fused with DNA replication proteins such as DnaB and RNA polymerase. These DNA replication proteins unwind the double-stranded DNA^12-14^, allowing cytosine deaminase to perform random deamination on the single-stranded DNA^15^, thereby introducing C→T mutations.

Currently, this random base editing tool has been constructed and applied in various chassis organisms such as *E. coli, S. cerevisiae*, and *Corynebacterium glutamicum* for whole-genome mutagenesis studies^16-19^. These tools have enabled the rapid acquisition of tolerant strains, such as ethanol-tolerant *C. glutamicum*, and cell chassis with high β-Carotene production in *S. cerevisiae*. Recently, Duan et al. utilized dockerin/cohesin-mediated protein−protein interactions to assemble DNA helicase, cytosine deaminase, and adenine deaminase in filamentous fungi^20^. Adenine deaminase TadA was originally an RNA-specific adenine deaminase. In 2017, an adenine deaminase called TadA-7.10 that acts on DNA adenine was developed using phage-assisted evolution technology^8^. However, TadA-7.10 exhibited low activity of deamination. In 2020, activity of the TadA enzyme was further improved through directed protein evolution, resulting in TadA-8e with eight additional mutations^21^. This variant demonstrated a 590-fold increase in enzyme activity compared to TadA-7.10. In 2023, researchers conducted further directed evolution on TadA to develop TadA variants with cytosine deaminase activity^22, 23^. As a result, the TadA-dual variant capable of simultaneously inducing A→G and C→T mutations was created. Nevertheless, Chen et al.^24^ and Wu et al.^25^ did not observe the anticipated dual-function editing phenotype in HEK293T cells and *Bacillus subtilis*, respectively. Recently, Fan et al. developed a base editor incorporating TadA-dual as the effector protein, which successfully achieved C→T and A→G mutations at target sites with exceptionally low off-target rates in rice and tomato cells^26^.

In this study, we investigated the mutations induced by TadA variants in whole-genome mutagenesis by fusing the RNA primase DnaG with TadA-8e and TadA-dual in *E. coli*. The DnaG-TadA fusion induced a high frequency of A→G mutations, while the DnaG-TadAD, when used in conjunction with UGI, introduced both C→T and A→G mutations in the *E. coli* genome. Both random editing tools significantly enhanced the rate of random mutations. The plasmids could eventually be cured to ensure the stability of the genome of the domesticated strain. We then applied these two tools to the adaptive evolution for Cd^2+^ and kanamycin tolerance, successfully achieving tolerant strains for 8.0 mM Cd^2+^ and 200 μg/mL kanamycin in just 17 d and 132 h, respectively. This work represents a significant advancement in the application of a single deaminase to simultaneously introduce C→T and A→G random mutations across an entire genome and is a newest attempt to explore rapid genomic mutational evolution using random base editing in *E. coli*.

**Table 1.** Strains used in this study. (The plasmids used in this study were Summarized in Table S1; speR: spectinomycin-resistant).

| Strain | Descriptions |
| --- | --- |
| *E. coli* DH5α | Used for plasmid construction |
| Eco-C | *E. coli* MG1655, pRSF1010-speR |
| Eco-DnaG-TadA | *E. coli* MG1655, pRSF1010-*dnaG-linker-(tadA-8e)*-speR |
| Eco-TadA-DnaG | *E. coli* MG1655, pRSF1010-*(tadA-8e)-linker-dnaG*-speR |
| Eco-DnaG-TadAD | *E. coli* MG1655, pRSF1010-*dnaG-linker-(tadA-dual)*-speR |
| Eco-DnaG-TadAD-UGI | *E. coli* MG1655, pRSF1010-*dnaG-linker-(tadA-dual)-ugi*-speR |
| Eco-DnaG-TadA-10G | *E. coli* MG1655, pRSF1010-*dnaG-linker-(tadA-8e)*-speR, Induction culture for ten generations |
| Eco-DnaG-TadAD-10G | *E. coli* MG1655, pRSF1010-*dnaG-linker-(tadA-dual)*-speR, Induction culture for ten generations |
| Eco-DnaG-TadAD-UGI-10G | *E. coli* MG1655, pRSF1010-*dnaG-linker-(tadA-dual)-ugi*-speR, Induction culture for ten generations |
| Eco-C-Cd | *E. coli* MG1655, pRSF1010-speR, enable to tolerate 2.5 mM Cd<sup>2+</sup> |
| Eco-DnaG-TadA-Cd | *E. coli* MG1655, pRSF1010-*dnaG-linker-(tadA-8e)*-speR, enable to tolerate 8.0 mM Cd<sup>2+</sup> |
| Eco-C-Km | *E. coli* MG1655, pRSF1010-speR, enable to tolerate 75 µg/mL kanamycin |
| Eco-DnaG-TadAD-UGI-Km | *E. coli* MG1655, pRSF1010-*dnaG-linker-(tadA-dual)-ugi*-speR, enable to tolerate 200 µg/mL kanamycin |

## RESULTS AND DISCUSSION

### Construction of a TadA-Mediated Random Base Editor in *E. coli*

The substrate for TadA-8e is adenine on single-stranded DNA, previous studies have attempted to increase genome mutation rates by expressing cytosine deaminase or adenine deaminase alone. However, deaminases without ssDNA did not increase the mutation rate significantly^16, 19^. As shown in **Figure 1**, we fused the RNA primase DnaG with TadA, aiming to randomly deaminate adenine to inosine (A→I) on the single-stranded DNA created by the RNA primase. Through subsequent complementary base pairing of the cell’s DNA, this process introduces random A→G mutations in the genome. We utilized a tandem inducible switch comprised of the LacI-based promoter and theophylline-responsive riboswitch to minimize leakage expression caused random mutation (**Figure 2A**). To explore the effects of spatial position on editing efficiency, we compared the fusion of TadA to either the N- or C-terminus of DnaG in *E. coli* MG1655 (**Figure 2B**) and characterized random mutations in the genome using rifampin resistance. Normally, rifampin inhibits the activity of the RNA polymerase β subunit, *rpoB*, leading to inhibited bacterial growth. When mutations occur in the amino acids (Q510, L511, Q510, Q513, F514, D516, H526, R529, S531, L533, G534) associated with rifampin binding to RpoB^27^, its inhibition ability is diminished. Thus, we used the number of rifampin-resistant colonies as a relative measure of the frequency of random mutations. As shown in **Figures 2C** and **2D**, the strain with TadA fused to the C-terminus of DnaG (hereafter referred to as Eco-DnaG-TadA) produced a significantly higher number of rifampin-resistant colonies, with an average of approximately 420 single colonies. In contrast, the strain with TadA fused to the N-terminus of DnaG (Eco-TadA-DnaG) yielded around 30 colonies rifampin-resistant colonies, representing a more than ten-folds difference between the two conditions. The N-terminus of DnaG has been found to bind to genomic DNA in *Mycobacterium tuberculosis*^14^. Fusing TadA to the N-terminus may similarly impair DnaG’s ability to bind to DNA effectively in *E. coli*, thereby reducing its capacity to unwind the DNA double helix. As expected, no rifampicin-resistant colonies were observed in the control strain (Eco-C). To characterize the random mutation frequency in the *E. coli* MG1655 without random base editor, we increased the plated bacterial density from OD_600_ of 0.01 to 0.1. In this case, the Eco-C produced an average of approximately 6 rifampicin-resistant colonies (**Figure S1**), it means that the mutation rates of Eco-DnaG-TadA increased by 664-fold. We randomly selected four single colonies and performed PCR and sequencing on the rifampin-resistance-related gene regions. The results revealed mutations in all the relevant regions, with all mutations being A→G (T→C) transitions (**Figure 2E**). One colony had a mutation that changed glutamine to arginine at position 513, while the other three colonies had mutations that converted histidine to arginine at position 526. These findings indicate that DnaG-TadA effectively increased the frequency of random mutations in the genome.

**Figure 1.**
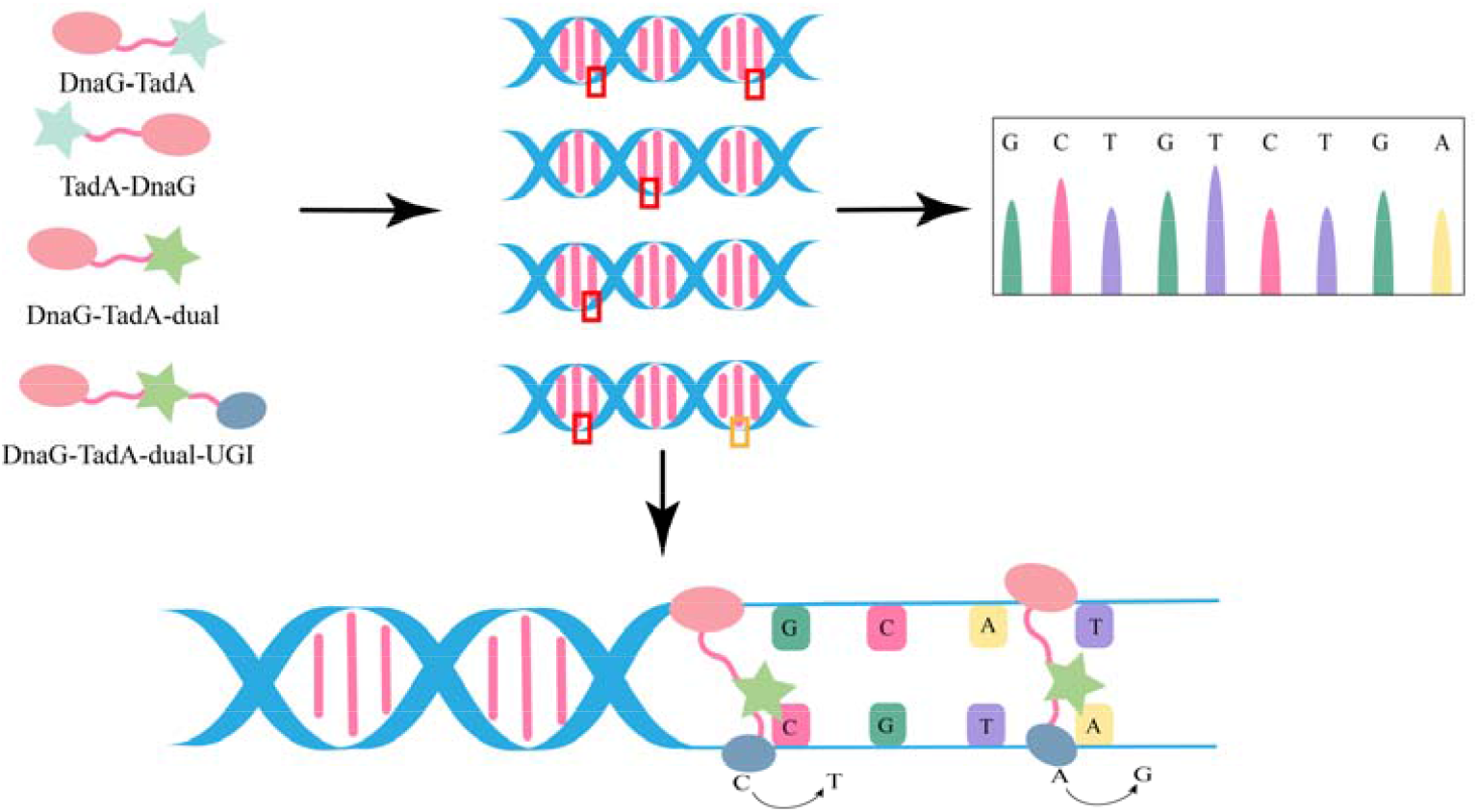
Schematic of random base editor guided by TadA variants in *E. coli*. The fusion expression of DnaG (RNA primase) and TadA variants introduces single A →G mutations or dual A→G and C→T mutations into the genome.

**Figure 2.**
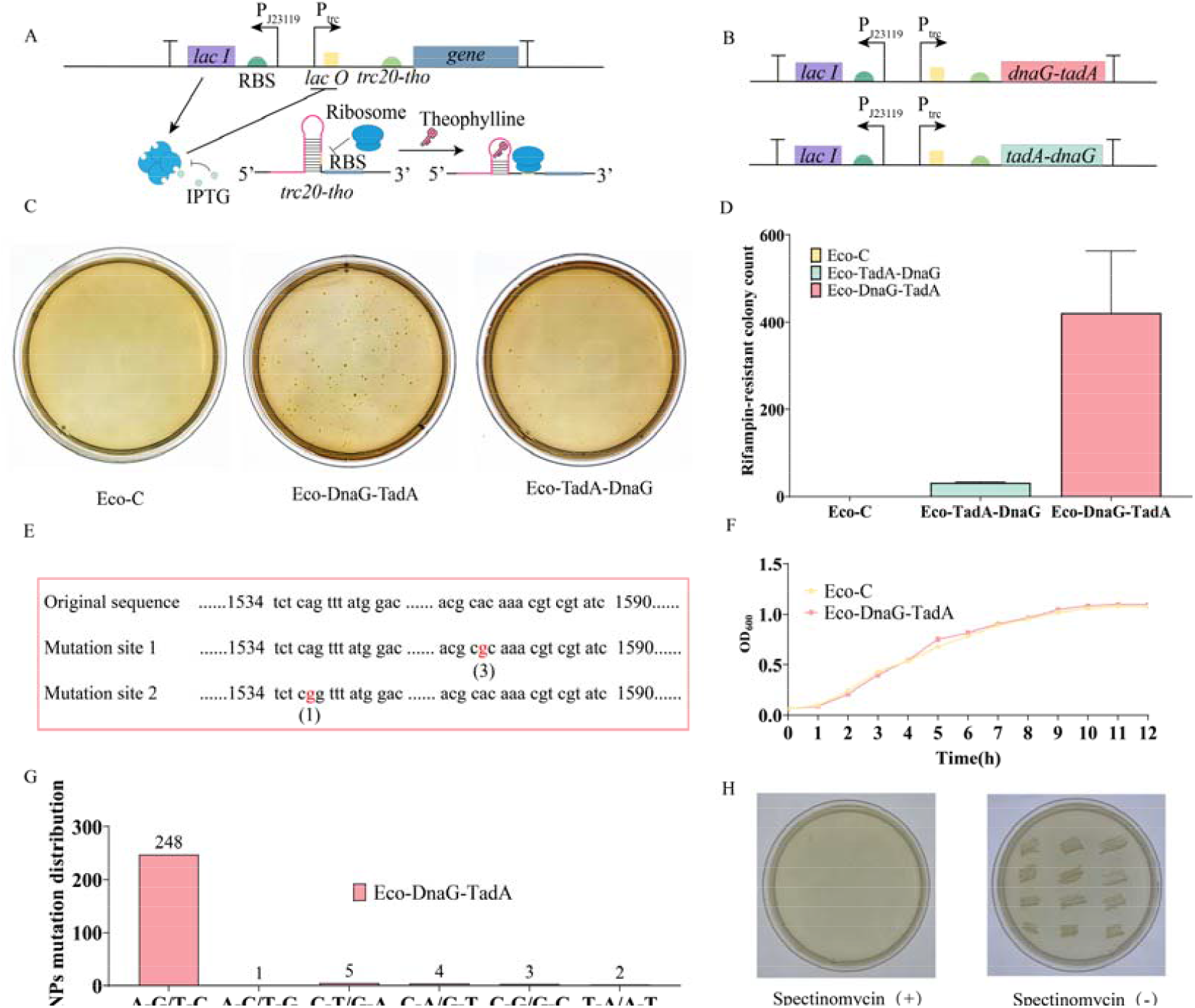
Construction and characterization of random base editing mediated by TadA in *E. coli*. (A) Schematic diagram of the tandem inducible switch. (B) Schematic of the engineered cassettes. (C) Pictures of single colonies from Eco-C, Eco-DnaG-TadA, and Eco-TadA-DnaG strains cultured overnight on rifampicin plates. (D) Statistical analysis of single colony numbers as mentioned above. (E) Sequencing analysis of *rpoB* gene mutations from selected rifampicin-resistant colonies of Eco-DnaG-TadA. (F) Growth comparison of Eco-C and Eco-DnaG-TadA. (G) Distribution of base mutation types in whole-genome sequencing of Eco-DnaG-TadA-10G. (H) Plasmid removal by using 10% SDS. Spectinomycin (+) meant medium with spectinomycin while Spectinomycin (-) meant medium without spectinomycin.

Next, we examined whether accelerated mutations into the genome via DnaG-TadA would affect cell growth. The results showed that the growth of the Eco-DnaG-TadA was comparable to that of the Eco-C (**Figure 2F**). This indicates that the expression of the fusion proteins in *E. coli* MG1655 does not cause severe growth inhibition, unlike tools reported previously that inhibit DNA mismatch repair genes which cause a decrease in growth^17^. The minimal impact on growth makes these fusion proteins particularly advantageous for applications such as adaptive evolution and related processes. To make sure the relevant plasmids can be easily removed from the strains once the desired mutations are achieved, we tested the plasmid cure strategy using 10% SDS for removing the pRSF1010-*dnaG-tadA* plasmid from Eco-DnaG-TadA. The results showed that 12 individual colonies selected using this method were not able to grow on media containing spectinomycin (**Figure 2H**), indicating that the plasmid had been successfully removed. To gain a more comprehensive understanding of the mutations induced by DnaG-TadA fusion protein in the genome, we cultivated the Eco-DnaG-TadA for 10 generations under induction and performed a whole-genome resequencing analysis (the according strain was Eco-DnaG-TadA-10G). A total of 263 single-base mutations were identified, with 248 being A→G (T→C) mutations in the Eco-DnaG-TadA-10G (**Figure 2G**). Additionally, there were 5 C→T (G→A) mutations. It is still unclear whether these mutations were resulted from the strain’s natural mutational processes or the previously reported cytidine deaminase activity of TadA-8e^28^. The occurrence of A→C (T→G), C→A (G→ T), C→G (G→C), and T→A (A→T) mutations was rare, each happening fewer than four times. These mutations may also be attributed to random deamination, where deaminated cytosine and adenine are excised by DNA glycosylases and incorrectly repaired by the cell’s mismatch repair system^29, 30^.

### Application of DnaG-TadA in Cd^2+^ Adaptation

We then applied DnaG-TadA in an adaptive laboratory evolution (ALE) using Cd^2+^ tolerance as a proof of concept. With 0.6 mM Cd^2+^ as the initial concentration, the resistant strains were transferred to medium with higher concentrations of Cd^2+^ after 24 hours’ cultivation. If the strains could not tolerate the respective concentration, they were re-cultivated at lower Cd^2+^ concentration. After 17 days’ cultivation, DnaG-TadA achieved a tolerance of 8.0 mM Cd^2+^, whereas the control strains only struggled to tolerate 2.5 mM Cd^2+^ (**Figure 3A**). Previously, Qin et al. reported using the GREACE method to accelerate the acclimation of cadmium-resistant *E. coli* strains, which took 2 months to reach 8.0 mM CdCl_2_^31^. Our ALE period was significantly shorter in comparison to the previous report. We isolated single colonies of the adapted strains (Eco-C-Cd and Eco-DnaG-TadA-Cd) and analyzed the growth of Eco-C-Cd and Eco-DnaG-TadA-Cd at 8.0 mM and 2.5 mM Cd^2+^ concentrations. The Eco-DnaG-TadA-Cd exhibited relatively suitable growth rates at both 2.5 mM and 8.0 mM Cd^2^□ concentrations (**Figure 3B**). In contrast, the Eco-C-Cd obtained from the control acclimation process was not able to grow at either concentration. Then, we conducted a whole-genome resequencing analysis of Eco-C-Cd and Eco-DnaG-TadA-Cd. Over a 17-days’ adaptive evolution period, the Eco-C-Cd produced only 4 base mutations, one of which was synonymous (**Figure 3C, 3D**). In contrast, the Eco-DnaG-TadA-Cd generated a total of 443 mutations in the same timeframe, including 241 non-synonymous mutations, 118 synonymous mutations, and 32 mutations in non-coding regions. Among them, the *zntA* gene, previously reported to be associated with Cd^2+^ resistance in *E. coli*^*32*^, mutated in both the Eco-C-Cd and Eco-DnaG-TadA-Cd, they all have the same M205I amino acid mutation site. In addition, 13 membrane protein-related genes or genes upstream of Eco-DnaG-TadA-Cd have also exhibited point mutations, and all of which are A→G (T→C). The membrane proteins are known to be involved in the Cd^2+^ adsorption^33^. In Eco-DnaG-TadA-Cd, an additional 74 mutations are associated with various transport proteins, the mutation type for all 74 mutations is consistently A→G (T→C). Among these transporters, CorA and ZitB are metal ions transport proteins, their mutations may be linked to Cd^2+^ tolerance^34, 35^. In addition, 11 other genes have been annotated as efflux transporter protein, some of these genes could be involved in the transport of metal ions out of cells^36, 37^.

**Figure 3.**
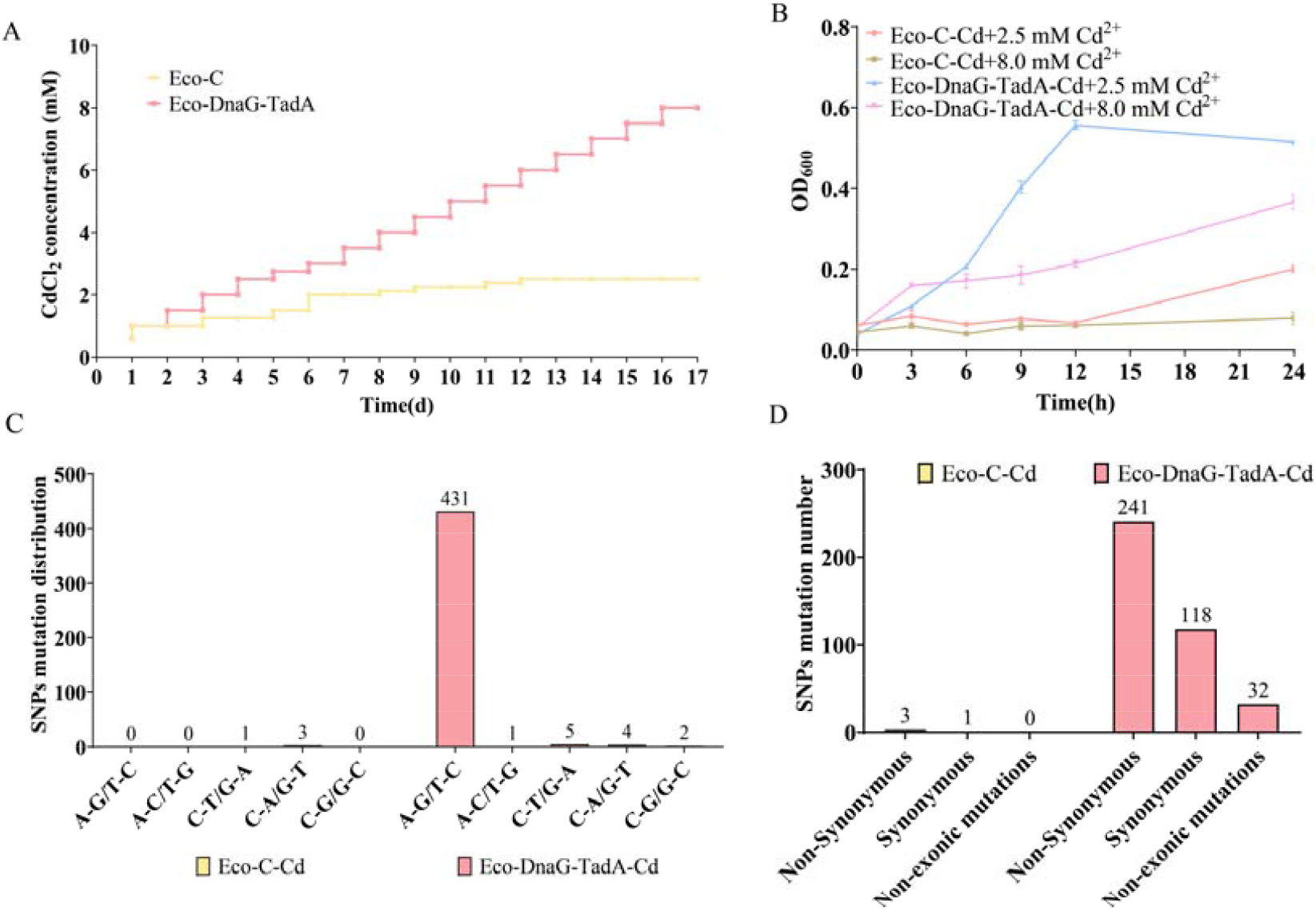
Accelerated adaptive evolution to Cd^2+^ by DnaG-TadA. (A) Conclusion of the Cd^2+^ evolutionary process for Eco-C and Eco-DnaG-TadA. (B) Growth of Eco-C-Cd and Eco-DnaG-TadA-Cd acclimated strains under 2.5 mM and 8.0 mM Cd^2+^ concentrations. (C) Distribution of base mutation types in whole-genome sequencing of Eco-C-Cd and Eco-DnaG-TadA-Cd acclimated strains. (D) Classification of point mutations in whole-genome sequencing.

### Construction and Evaluation of the Dual Base Editor

To further extend the base editing libraries, we replaced TadA-8e with the a mutant TadA-dual^22^, which has been reported with dual-function editing capabilities, resulting in the construction of Eco-DnaG-TadAD (**Figure 4A**). Additionally, the UGI (uracil glycosylase inhibitor) derived from *Bacillus subtilis* phage has been shown to effectively inhibit UDG (uracil-DNA glycosylase) in *E. coli* and other species^38, 39^, thereby reducing the mismatch repair of U formed by cytosine deamination and enhancing C→T editing efficiency. Therefore, we fused UGI to the C-terminus of TadA to potentially increase the frequency of C→T mutations in the random dual base editor. Similarly, we used rifampicin resistance to characterize the random mutation frequency. Both produced abundant rifampicin-resistant colonies, with Eco-DnaG-TadAD generating an average of 196 colonies and Eco-DnaG-TadAD-UGI producing approximately 234 colonies on average (**Figure 4B, 4C**). The mutation rates of Eco-DnaG-TadAD and Eco-DnaG-TadAD-UGI were increased by 309 and 370-fold, respectively. However, the mutation rates were significantly decreased compared to Eco-DnaG-TadA. This decrease might be due to the fact that TadA sacrificing some of its enzymatic activity during the directed evolution for dual-function mutations, in previous research by David Liu’s team, the TadA-dual-guided base editor (TadDE) exhibited relatively lower adenine deamination activity at certain editing sites compared to the TadA-8e-guided ABE8e base editor^22^. We evaluated the growth curves of both Eco-DnaG-TadAD and Eco-DnaG-TadAD-UGI (**Figure 4D**), and then compared to the Eco-C (**Figure 2F**), but no growth inhibition was observed.

**Figure 4.**
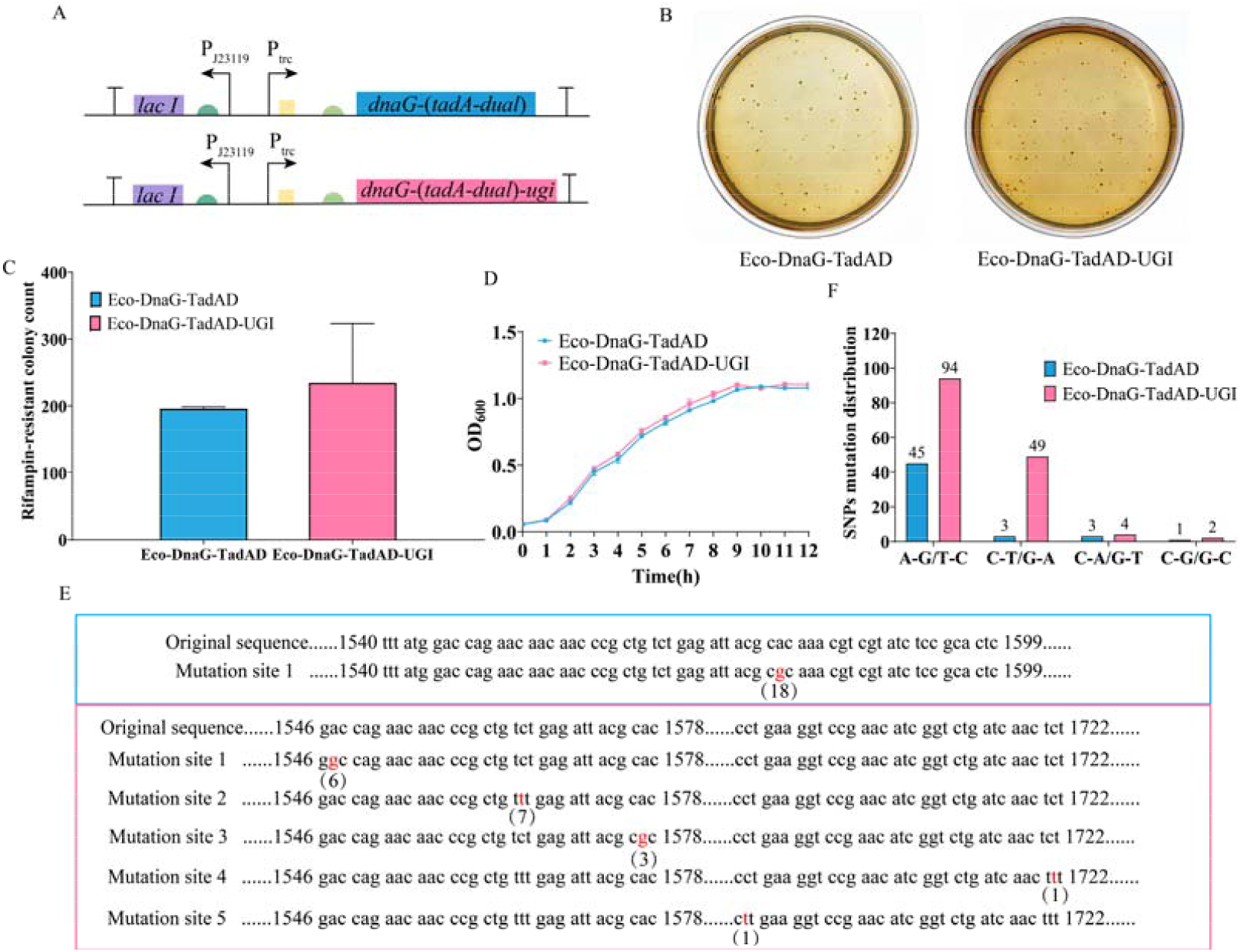
Construction and characterization of random dual base editing mediated by TadA-dual in *E. coli*. (A) Schematic representation of the engineered cassettes. (B) Pictures of single colonies from Eco-DnaG-TadAD, and Eco-DnaG-TadAD-UGI strains cultured on rifampicin plates. (C) Statistical analysis of single colony numbers. (D) Growth curve of the Eco-DnaG-TadAD and Eco-DnaG-TadAD-UGI. (E) Sequencing analysis of *ropB* gene mutations from selected rifampicin-resistant colonies, the sequencing results of Eco-DnaG-TadAD are in the blue box, and the sequencing results of Eco-DnaG-TadAD-UGI are in the pink box. (F) Distribution of base mutation types in whole-genome sequencing of Eco-DnaG-TadAD and Eco-DnaG-TadAD-UGI.

To clarify the mutation types, we selected 18 rifampicin-resistant colonies each from the Eco-DnaG-TadAD and Eco-DnaG-TadAD-UGI mutants for sequencing analysis of the *rpoB* region (**Figure 4E**). All 36 colonies exhibited mutations at or near the rifampicin resistance-related amino acid sites, indicating that the observed resistance was mutation-induced. Notably, all 18 Eco-DnaG-TadAD colonies had only A→G mutations in the corresponding gene region, especially exhibiting a mutation of H526R. Among the 18 Eco-DnaG-TadAD-UGI colonies, 9 exhibited A→G (T→C) mutations. Six of these had D516G mutations, changing the codon from GAC to GGC, while three had H526R mutations, changing the codon from CAC to CGC. The remaining 9 colonies showed C→T (G→A) mutations, with seven occurring at position 522, changing serine (TCT) to phenylalanine (TTT). The other two mutations were P564L and S574F. These results suggested that, under the cellular mismatch repair mechanism relying on UDG, TadA-dual produces C→T (G→A) mutations at a much lower frequency compared to A→G (T→C) mutations. However, when UDG’s activity was inhibited by UGI, the Eco-DnaG-TadAD-UGI achieved a comparable mutation rate for both C→T (G→A) and A→G (T→C) mutations.

Gene sequencing of rifampicin-resistant colony did not detect C→T (G→A) mutation activity for Eco-DnaG-TadAD initially, possibly due to an insufficient number of samples. Therefore, we performed ten successive rounds of induction on Eco-DnaG-TadAD and Eco-DnaG-TadAD-UGI and conducted a whole-genome sequencing analysis to investigate their dual-random base editing activity. As shown in **Figure 4F**, Eco-DnaG-TadAD-10G and Eco-DnaG-TadAD-UGI-10G introduced 52 and 149 mutations, respectively. The mutation number was decreased compared to Eco-DnaG-TadA-10G, consistent with the reduction in the number of rifampicin-resistant colonies. Furthermore, Eco-DnaG-TadAD primarily generated A →G (T→C) mutations, with a total of 45 occurrences, indicating that TadA-dual, when expressed alone with DnaG, cannot introduce a wide variety of high-frequency mutations into the genome. In contrast, among the mutations produced by Eco-DnaG-TadAD-UGI, there were 94 A→G (T→C) mutations and 49 C→T (G→A) mutations, demonstrating that this tool has the significant ability to introduce 2 types of base mutations into the genome.

### Application of DnaG-TadAD-UGI in Kanamycin Adaptation

Finally, we applied DnaG-TadAD-UGI to the adaptive evolution towards kanamycin. Remarkably, within just 132 h, DnaG-TadAD-UGI enabled the strain to tolerate 200 μg/mL of kanamycin, which is much higher than the control strain (75 μg/mL) (**Figure 5A**). Meanwhile, the Eco-DnaG-TadAD-UGI-Km exhibited rapid growth at both 75 μg/mL and 200 μg/mL kanamycin concentrations, whereas the Eco-C showed a much slower growth (**Figure 5B**). This indicates DnaG-TadAD-UGI substantially enhances the strain’s ability to rapidly generate mutations, leading to branches that can withstand adverse conditions. Although its mutation rate is lower than that of DnaG-TadA, the DnaG-TadAD-UGI can produce a wider variety of mutations. This diversity might result in more potential mutations, suggesting that DnaG-TadAD-UGI and DnaG-TadA each have unique advantages for practical applications.

**Figure 5.**
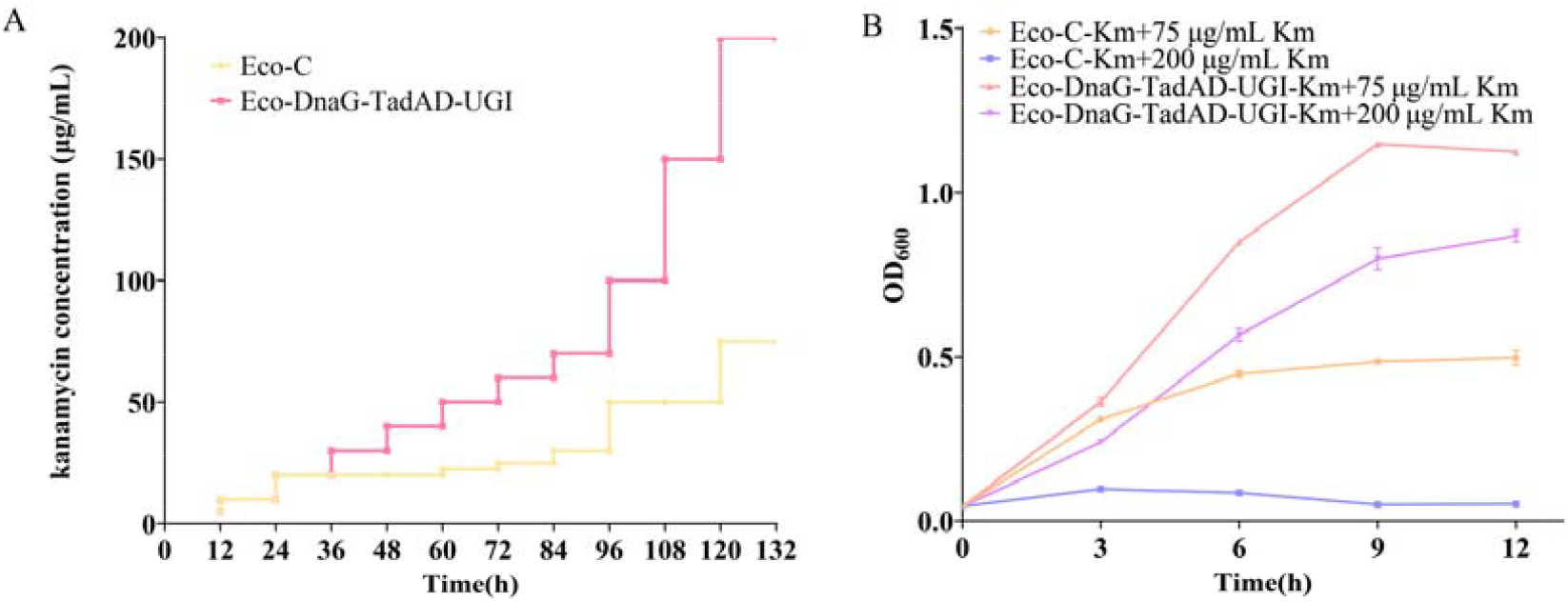
Adaptive evolution towards tolerance of kanamycin by DnaG-TadAD-UGI. (A) Conclusion of the kanamycin adaptive process for control and DnaG-TadAD-UGI. (B) Growth of control and DnaG-TadAD-UGI kanamycin acclimated strains under 75 μg/mL and 200 μg/mL of kanamycin concentrations.

## METHODS

### Strain and Plasmid Construction

*E. coli* DH5α was used for plasmid construction. Gene information is provided in the supplementary **Table S1**. The DnaG gene was sourced from the *E. coli* genome, the gene sequence of the linker and TadA-8e from the HMGN1-A8e ^40^. The plasmid backbone was pRSF1010^41^. All PCR reactions were carried out using high-fidelity DNA polymerase (Vazyme, Nanjing, China), and all plasmids were assembled via Gibson assembly (Vazyme, Nanjing, China). Sequencing analysis was performed by Genewiz (Suzhou). The constructed plasmids were chemically transformed into *E. coli* MG1655. Both *E. coli* strains were cultured at 37°C with a shaking speed of 160 rpm (HNYC202T, Illuminating Shaker, Honour, Tianjin, China).

### Cultivation and Growth Testing of Evolved Cells

*E. coli* strains were cultivated overnight at 37°C in LB medium. The OD_600_ of the cultures was measured using a microplate reader (ALLSHENG, AMR-100, Hangzhou, China). The cultures were then inoculated into fresh medium, adjusting the OD_600_ to 0.04 by varying the inoculum volume. Additionally, 0.5 mM IPTG and 2 mM theophylline, along with 100 μg/mL spectinomycin, were added to the medium. To monitor the growth of the strains, OD_600_ was measured every three hours, and the growth was characterized based on changes in OD_600_.

### The Plasmid Curing by 10% SDS

The plasmid curing method follows procedures outlined in relevant literature^42^. The concentration of the bacterial strain was diluted to approximately 10^4 CFU/mL using fresh LB liquid medium. A 0.5 mL aliquot of the diluted bacterial culture was added to 4.5 mL of LB liquid medium containing 10% SDS. The mixture was incubated at 37°C with shaking at 160 rpm for 48 hours. After incubation, the cultures were centrifuged to remove the supernatant and the pellet was washed with fresh LB medium to remove residual SDS. The cultures were then serially diluted and plated on fresh LB agar plates, followed by incubation for 24 hours. Single colonies were picked and streaked onto LB agar plates with and without spectinomycin. These plates were incubated at 37°C for 12 hours. The growth differences on the two media were observed: colonies that grew on the antibiotic-free plates but not on the spectinomycin plates were identified as successfully cured of the plasmid.

### Rifampicin Assay

Under the specified culture conditions and with the addition of inducers, cells were induced for 12 hours. Then, 0.01 OD_600_ of the culture was plated onto solid media containing 50 μg/mL rifampicin. The plates were incubated at 37°C for 20 hours, and the number of colonies was counted. Each test strain was set up with three biological replicates. The primers RpoB-F: actggtagaaatctaccgcatg and RpoB-R: cagagacacacacggcatct were used for PCR amplification of the *rpoB* gene region. The sequencing reactions were performed and analyzed by GENEWIZ (Suzhou).

### Adaptive Evolution towards Cd^2+^

The *E. coli* MG1655 carrying the control plasmid or DnaG-TadA plasmid were cultured in fresh LB medium, then pre-culture them in LB medium containing inducers with 0.6% CdCl_2_ concentration. Incubate at 37°C with shaking at 160 rpm for 24 hours. When the OD_600_ of the strains exceeds 0.2, wash the cells with fresh medium and then inoculate them into LB medium containing inducers with a higher concentration of CdCl_2_.

### Adaptive Evolution towards Antibiotics

The *E. coli* MG1655 carrying the control plasmid or DnaG-TadAD-UGI plasmid were cultured in fresh LB medium, then pre-culture them in LB medium containing inducers with an initial antibiotic concentration of 5μg/mL. Incubate at 37°C with shaking at 160 rpm for 12 hours. When the OD_600_ of the strains exceeds 0.2, wash the cells with fresh medium and then inoculate them into LB medium containing inducers with a higher concentration of antibiotics.

### Whole Genome Sequencing

Single colonies of DnaG-TadA, DnaG-TadAD, and DnaG-TadAD-UGI after ten generations of induction, as well as single colonies of DnaG-TadA-8.0 mM Cd^2+^-resistant strains and control-2.5 mM Cd^2+^-resistant strains were isolated. Genomic DNA was extracted using the Magen Hipure Soil DNA Kit. Sequencing libraries were prepared using the standard library construction procedure of the VAHTS Universal Plus DNA Library Prep Kit for Illumina V2. Sequencing was performed on the Illumina NovaSeq PE150 platform by GENEWIZ (Suzhou), with a sequencing depth of approximately 150 and filtered reads of about 2G.

## ACKNOWLEDGMENTS

This research was supported by grants from the National Key Research and Development Program of China (Grant no. 2019YFA0904600), the National Natural Science Foundation of China (Grant nos. 32371486 and 32270091).

## Author Contributions

H. W. performed the experiments and wrote the manuscript; H. W., Z. D. and J. S. analyzed the data; L. C., T. S. and W. Z. designed the study and revised the manuscript.

### Notes

All the authors declare no competing financial interest.

## Notes

### Competing Interest Statement

The authors have declared no competing interest.

