## Supplemental Tables and figures for "Genome-wide A→G and C→T Mutations Induced by Functional TadA Variants in *Escherichia coli*"

**Supplementary Figure S1.**

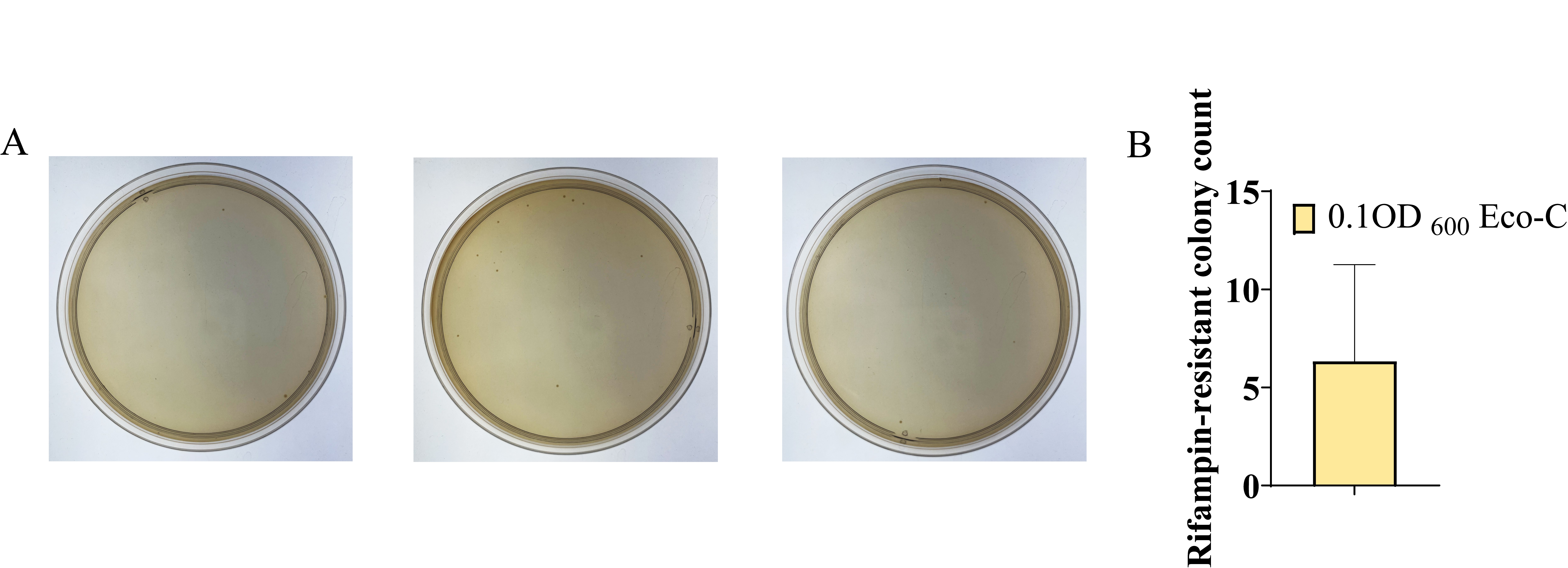

**Supplementary Figure S1.** (A) Pictures of single colonies from 0.1OD_600_ Eco-C cultured on rifampicin plates. (B) Statistical analysis of single colony numbers.

**Supplementary Table S1.** The plasmids used in this study.

| **Plasmid** | **Description** |
| --- | --- |
| pRSF1010-speR | speR; *E. coli* episomal plasmids |
| pRSF1010-*dnaG*-*linker*-(*tadA-8e*)-speR | pRSF1010 derivative harboring *dnaG*-*linker*-(*tadA-8e*), under the control of *lacI* and theophylline riboswitch, for expression of *dnaG*-*linker*-(*tadA-8e*). |
| pRSF1010-(*tadA-8e*)-*linker*-*dnaG*-speR | pRSF1010 derivative harboring (*tadA-8e*)-*linker*-*dnaG*, under the control of *lacI* and theophylline riboswitch, for expression of (*tadA-8e*)-*linker*-*dnaG*. |
| pRSF1010-*dnaG*-*linker*-(*tadA-dual*)-speR | pRSF1010 derivative harboring *dnaG*-*linker*-(*tadA-dual*), under the control of *lacI* and theophylline riboswitch, for expression of *dnaG*-*linker*-(*tadA-dual*). |
| pRSF1010-*dnaG*-*linker*-(*tadA-dual*)-*ugi*-speR | pRSF1010 derivative harboring *dnaG*-*linker*-(*tadA-dual*)-*ugi*, under the control of *lacI* and theophylline riboswitch, for expression of *dnaG*-*linker*-(*tadA-dual*)-*ugi*. |

**Supplementary Table S2.** The genes used in this study.

| **Gene** | **Sequence(5’→3’)** |
| --- | --- |
| *dnaG* | atggctggacgaatcccacgcgtattcattaatgatctgctggcacgcactgacatcgtcgatctgatcgatgcccgtgtgaagctgaaaaagcagggcaagaatttccacgcgtgttgtccattccacaacgagaaaaccccgtccttcaccgttaacggtgagaaacagttttaccactgctttggatgtggcgcgcacggcaacgcgatcgacttcctgatgaactacgacaagctcgagttcgtcgaaacggtcgaagagctggcagcaatgcacaatcttgaagtgccatttgaagcaggcagcggccccagccagatcgagcgccatcagaggcaaacgctttatcagttgatggacggtctgaatacgttttaccaacaatctttacaacaacctgttgccacgtctgcgcgccagtatctggaaaaacgcggattaagccacgaggttatcgctcgctttgcgattggttttgcgccccccggctgggacaacgtcctgaagcggtttggcggcaatccagaaaatcgccagtcattgattgatgcggggatgttggtcactaacgatcagggacgcagttacgatcgtttccgcgagcgggtgatgttccccattcgcgataaacgcggtcgggtgattggttttggcgggcgcgtgctgggcaacgatacccccaaatacctgaactcgccggaaacagacattttccataaaggccgccagctttacggtctttatgaagcgcagcaggataacgctgaacccaatcgtctgcttgtggtcgaaggctatatggacgtggtggcgctggcgcaatacggcattaattacgccgttgcgtcgttaggtacgtcaaccaccgccgatcacatacaactgttgttccgcgcgaccaacaatgtcatttgctgttatgacggcgaccgtgcaggccgcgatgccgcctggcgagcgctggaaacggcgctgccttacatgacagacggccgtcagctacgctttatgtttttgcctgatggcgaagaccctgacacgctagtacgaaaagaaggtaaagaagcgtttgaagcgcggatggagcaggcgatgccactctccgcatttctgtttaacagtctgatgccgcaagttgatctgagtacccctgacgggcgcgcacgtttgagtacgctggcactaccattgatatcgcaagtgccgggcgaaacgctgcgaatatatcttcgtcaggaattaggcaacaaattaggcatacttgatgacagccagcttgaacgattaatgccaaaagcggcagagagcggcgtttctcgccctgttccgcagctaaaacgcacgaccatgcgtatacttatagggttgctggtgcaaaatccagaattagcgacgttggtcccgccgcttgagaatctggatgaaaataagctccctggacttggcttattcagagaactggtcaacacttgtctctcccagccaggtctgaccaccgggcaacttttagagcactatcgtggtacaaataatgctgccacccttgaaaaactgtcgatgtgggacgatatagcagataagaatattgctgagcaaaccttcaccgactcactcaaccatatgtttgattcgctgcttgaactgcgccaggaagagttaatcgctcgtgagcgcacgcatggtttaagcaacgaagaacgcctggagctctggacattaaaccaggagctggcgaaaaag |
| *Linker1*(between *tadA* variant and *dnaG*) | tctggaggatctagcggaggatcctctggaagcgagacaccaggcacaagcgagtccgccacaccagagagctccggcggctcctccggaggatcc |
| *tadA-8e* | atgtccgaagtggaattctcccacgaatactggatgcgccacgcactgaccctggcaaagcgcgcacgcgatgaacgcgaagtgcctgtgggcgcagtgctggtgctgaacaaccgcgtgatcggcgaaggctggaaccgcgcaatcggcctgcacgatccaaccgcacacgcagaaatcatggcactgcgccaaggcggcctggtgatgcagaactaccgcctgatcgatgcaaccctgtacgtgaccttcgaaccatgcgtgatgtgcgccggcgcaatgatccactcccgcatcggccgcgtggtgttcggctggcgcaactccaagcgcggcgcagccggctccctgatgaacgtgctgaactaccctggcatgaaccaccgcgtggaaatcaccgaaggcatcctggcagatgaatgcgcagcactgctgtgcgatttctaccgcatgccacgccaagtgttcaacgcacagaagaaggcacagtcctccatcaac |
| *tadA-dual* | atgtccgaagtggaattctcccacgaatactggatgcgccacgcactgaccctggcaaagcgcgcacgcgatgaaGGAgaaGCGcctgtgggcgcagtgctggtgctgaacaaccgcgtgatcggcgaaggctggaaccgcCGTatcggcctgcacgatccaaccgcacacgcagaaatcatggcactgcgccaaggcggcctggtgatgcagaacTCCcgcctgatcgatgcaaccctgtacgtgaccttcgaaccatgcgtgatgtgcgccggcgcaatgatcAACtcccgcatcggccgcgtggtgttcggcGTGcgcaactccaagcgcggcgcagccggctccctgatgaacgtgctgaactaccctggcatgaaccaccgcgtggaaatcaccgaaggcatcctggcagatgaatgcgcagcactgctgtgcgatttctaccgcatgccacgccaagtgttcaacgcacagaagaaggcacagtcctccatcaac |
| *linker2*(between *tadA-dual* and *ugi*) | tctggtggaagcggaggatctggcggcagc |
| *ugi* | atgacgaatctcagcgacatcatcgagaaagaaacggggaaacaactggtgatccaggaaagcatcctgatgctccccgaggaggtcgaggaagtcatcggcaataaaccggagtcggatatcctggtccataccgcgtatgacgaaagcacggacgagaacgtcatgctcctcacgtcggatgcgccggaatataaaccgtgggccctcgtgatccaagatagcaatggcgagaacaaaatcaagatgctc |
